# Semi-blind sparse affine spectral unmixing of autofluorescence-contaminated micrographs

**DOI:** 10.1101/529008

**Authors:** Blair J. Rossetti, Steven A. Wilbert, Jessica L. Mark Welch, Gary G. Borisy, James G. Nagy

## Abstract

Spectral unmixing methods attempt to determine the concentrations of different fluorophores present at each pixel location in an image by analyzing a set of measured emission spectra. Unmixing algorithms have shown great promise for applications where samples contain many fluorescent labels; however, existing methods perform poorly when confronted with autofluorescence-contaminated images. We propose an unmixing algorithm designed to separate fluorophores with overlapping emission spectra from contamination by autofluorescence and background fluorescence. First, we formally define a generalization of the linear mixing model, called the affine mixture model (AMM), that specifically accounts for background fluorescence. Second, we use the AMM to derive an affine nonnegative matrix factorization method for estimating fluorophore endmember spectra from reference images. Lastly, we propose a semi-blind sparse affine spectral unmixing (SSASU) algorithm that uses knowledge of the estimated endmembers to learn the autofluorescence and background fluorescence spectra on a per-image basis. When unmixing real-world spectral images contaminated by autofluorescence, SSASU greatly improved proportion indeterminacy as compared to existing methods for a given relative reconstruction error. The source code used for this paper was written in Julia and is available with the test data at https://github.com/brossetti/ssasu.

## I. INTRODUCTION

Most conventional fluorescence microscopes use a series of mirrors and optical filters to separate the emission light of different fluorophores. The characteristics of these filters (i.e., what wavelengths of light they let pass) dictate what types of fluorophores and, more importantly, how many fluorophores can be used in an experiment. There exist a number of tools to help optimize the choice of fluorophores based on the configuration of a given microscope, such as the recently released SPEKcheck by Phillips et al. [2018]. These tools attempt to reveal the set of fluorophores that minimize spectral cross talk, a problem where the filter used for one fluorophore does not adequately exclude the fluorescent emission of other fluorophores (see Waters [2009] and references therein). Yet, even with such optimization, most experiments are still limited to three or four fluorescent labels. To make matters worse, many biological samples contain a variety of autofluorescent molecules that emit light in wavelengths that overlap and obscure the desired signal.

Spectral microscopy has become the method of choice when needing to avoid cross talk, mitigate autofluorescence, and simultaneously visualize many biological objects [Harmany et al., 2017, Jonkman et al., 2014, Levenson et al., 2008, Valm et al., 2011, 2016]. Instead of relying on filters, spectral microscopes use specialized optics and computational analysis to measure and separate the light emitted by different fluorophores. A spectral image is a three-dimensional data structure that holds a discrete spectral emission profile for each (*x, y*) pixel location. The range of detectable wavelengths and the number and width of the spectral bands varies by system and application. However, most commercial spectral microscopes are capable of measuring tens (multispectral) or even hundreds (hyperspectral) of spectral bands at resolutions of 1-15 nm/band [Cole et al., 2013, Gao and Smith, 2015]. For an in-depth treatment of spectral microscopy and its applications, we refer the reader to the “Spectral Imaging” special issue of *Cytometry Part A* [Lerner et al., 2006] and the reviews by Garini and Tauber [2013] and Li et al. [2013].

Since a spectral microscope simply records the spectrum at each pixel, the ability to separate fluorophores with overlapping emission spectra depends on the choice of spectral unmixing algorithm. Spectral unmixing refers to a group of techniques that attempt to determine how much of each fluorophore was present in some observed spectrum. Nearly all of these methods assume that light emitted by different fluorophores mix linearly, and are rightly called linear mixing models (LMM) [Zimmermann et al., 2014]. In the deterministic case, fluorophores are assumed to have one canonical emission spectrum called a *reference spectrum* or *endmember* (a term borrowed from mineralogy that refers to the purest form of an element that exists in a mixture). We can represent the set of *N* endmembers, each with *M* spectral bands, as the columns, denoted **s**_*n*_ for *n* = 1, …, *N*, of the endmember matrix 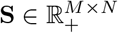. If we denote the endmember concentrations, or weights, as 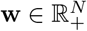, then we can write the LMM as

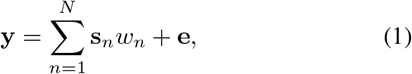

where 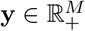 is the observed spectrum, *w*_*n*_ is the *n*^*th*^ entry of **w**, and **e** is i.i.d. noise (typically assumed to be Gaussian or Poisson distributed). When considering an entire spectral image with *P* pixels, denoted 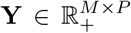, we can rewrite the LMM in matrix form as

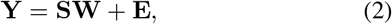

where **E** ∈ ℝ_*M×P*_ is noise and 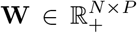 is the set of weights for each endmember in each pixel. It is important to highlight that the entries of **W** must be nonnegative (i.e., in ℝ_+_) since negative combinations of endmembers are not physically meaningful. The endmember weights are visualized by reorganizing the columns of **W** to produce a three-dimensional (*P*_*y*_ × *P*_*x*_ × *N*) unmixed image, where *P*_*y*_ and *P*_*x*_ are the number of vertical and horizontal pixels, respectively.

When the number and spectra of endmembers in an image are known in advance, spectral unmixing under the LMM is equivalent to the problem of Nonnegative Least Squares (NLS), and it can be written as

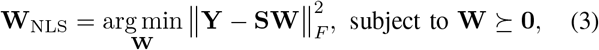

where and ‖·‖_*F*_ is the Frobenius norm, ⪰ is element-wise ≥, and **0** is an appropriately-sized matrix of zeros. Conveniently, NLS is a strictly convex optimization problem with a unique solution when the endmembers are linearly independent. Many unmixing algorithms that come packaged with commercial microscopes rely on some variant of the NLS active-set algorithm by Lawson and Hanson [1995]. While NLS is sufficient for some applications, it is not always possible to know which endmembers exist in an image. In particular, there are many different types of autofluorescent molecules, and knowing which autofluorescence endmembers are present is often impractical. In addition, endmembers are typically estimated from a reference sample. A poorly prepared reference sample or improper estimation method will lead to undesired unmixing results [Zimmermann et al., 2014]. As such, NLS lacks the flexibility to adequately handle many of the unmixing problems that arise in realistic applications.

Spectral unmixing methods have been used extensively by the remote sensing community for the analysis of hyperspectral geospatial data, and there exists a rich literature on advanced hyperspectral unmixing algorithms [Bioucas-Dias et al., 2012, Drumetz et al., 2016, Heylen et al., 2014, Keshava and Mustard, 2002, Keshava, 2003, Ma et al., 2014]. Although some methods from remote sensing have been directly applied to spectral micrographs [Harris, 2006, Lu and Fei, 2014], there are several key differences between the problem conditions that render many unmixing algorithms for geospatial data unsuitable for spectral microscopy. As compared to remote sensing data,

1. spectral micrographs contain fewer spectral bands and, therefore, suffer less from the curse of dimensionality;
2. spectral micrographs have significantly higher spatial resolution relative to the size of the target objects, meaning that individual pixels contain fewer endmembers and neighboring pixels have similar compositions;
3. the number of fluorophores used for labeling is known *a priori*;
4. it is possible to estimate endmembers from reference images;
5. and spectral micrographs are often heavily contaminated by background fluorescence and autofluorescent organic molecules.

As a result of these differences, there is an existing and ongoing need to develop spectral unmixing methods specifically tailored towards spectral microscopy applications [Arena et al., 2017].

Several notable algorithms have been recently introduced with the aim of unmixing images contaminated by autofluorescence. Fereidouni et al. [2012, 2014] developed a fast unmixing algorithm (along with an ImageJ plugin) inspired by the phasor methods used in lifetime imaging. Unfortunately, spectral phasor analysis is a geometric approach that is computationally expensive when unmixing more than three endmembers. A highly optimized phasor approach was proposed by Cutrale et al. [2017] for classifying autofluorescence-contaminated spectral images. Their Hyper-Spectral Phasors (HySP) software was able to differentiate four fluorophores and three autofluorescent sources in a single volumetric time-lapse data set. Unlike unmixing methods, however, HySP assigns each pixel to an endmember class and does not determine the relative proportions of each endmember. As a result, HySP is not suitable for experiments where fluorophores or autofluorescent molecules spatially overlap. Megjhani et al. [2017] proposed a powerful morphologically constrained spectral unmixing (MCSU) algorithm using dictionary learning. In addition to learning the endmember spectra from reference images, MCSU builds a dictionary of morphological motifs (e.g., cell or organelle shapes) unique to each fluorophore. Although MCSU shows impressive results for up to eight fluorophores, it requires that the reference images share the same morphologies found in the test images and that these morphologies differ between fluorophores.

The sparse unmixing by variable splitting and augmented Lagrangian (SUnSAL) algorithm by Bioucas-Dias and Figueiredo [2010] and its spatially regularized variant SUnSAL-TV by Iordache et al. [2012] are two useful methods for images containing endmembers that may not be linearly independent (recall that **S** must have full column rank for NLS to have a unique solution). The imposed sparsity and total variation (TV) regularizers help guide the optimization towards sparse solutions in which fewer endmembers are used to explain the observed spectral signatures. As with NLS, however, both SUnSAL and SUnSAL-TV are unable to learn or adjust endmembers on a per-image basis.

Perhaps the most popular open source unmixing tool is the PoissonNMF plugin for ImageJ by Neher et al. [2009]. PoissonNMF is based on the nonnegative matrix factorization (NMF) methods popularized by Lee and Seung [2001]. In the context of unmixing, NMF is a blind source separation algorithm that aims to learn the endmembers and their weights by solving

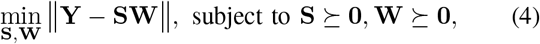

for an appropriate distance metric. As its name suggests, PoissonNMF is intended for use with images affected primarily by Poisson distributed noise (i.e., shot-noise). The PoissonNMF ImageJ plugin can operate in a blind or semi-blind manner where known endmembers can be fixed when solving Eq. 4. However, Neher et al. [2009] report that PoissonNMF yields unsatisfactory results when unmixing more than four endmembers due to the heavily overlapping emission profiles.

Similar semi-blind NMF approaches using the Gaussian noise model for read-noise-limited data have been suggested by Huang et al. [2015], Tong et al. [2016], and Qin et al. [2016]. These methods rely on sparsity regularization to help with linearly dependent endmembers, and they have been shown to work well on images with three to five endmembers. However, none of these methods separately and explicitly model contamination by autofluorescence from organic molecules and background fluorescence from ambient and stray light.

Global background fluorescence originates from a variety of sources in the sample and optical path [Waters, 2009]. When neglecting this source of light, the weight matrix becomes dense and sparsity constraints are ineffective. This occurs because the offset intensity stemming from background fluorescence violates the assumed zero-mean Gaussian noise model. Laurberg and Hansen [2007] introduced a sparse affine NMF method aimed at handling the offset components found in a variety of data types. Woolfe et al. [2011] used a similar affine model (incorrectly called a linear model) to address autofluorescence in spectral images; however, this model did not impose constraints on sparsity.

In this paper, we formally define an affine mixing model (AMM) that generalizes the LMM by including a term to absorb any background fluorescence. From this model, we derive an affine nonnegative matrix factorization (ANMF) method for estimating endmembers from reference images. We then propose a semi-blind sparse affine spectral unmixing (SSASU) method for images contaminated with both autofluorescence and background fluorescence.

We test the proposed methods on the study of complex bacterial biofilms. As shown by Valm et al. [2011] and Mark Welch et al. [2016], multiplexed labeling and spectral microscopy can be used to explore the spatial relationships within biofilms. In these experiments, fluorophores conjugated to oligonucleotide probes were used to label and differentiate tens of bacterial taxons. Until now, the presence of autofluorescence and background has made separating these overlapping fluorophores difficult for certain biofilm samples. We show that our SSASU approach is able to successfully reduce the impact of autofluorescence and background and unmix seven fluorophores.

## II. MATERIALS AND METHODS

### A. Sample preparation

To evaluate the proposed method, a set of seven reference samples, ten test samples, and one no-probe control was prepared. The bacteria *Leptotrichia buccalis* was used as the biological target for generating reference samples for each of the seven fluorophores: DY-415, DY-490, ATTO 520, ATTO 550, Texas Red-X, ATTO 620, and ATTO 655 (Dyomics GmbH; ATTO-TEC GmbH; Thermo Fisher Scientific Inc.). *L. buccalis* cells were cultured, fixed, and then separately hybridized using custom fluorophore-conjugated oligonucleotide probes (biomers.net GmbH) as described by Mark Welch et al. [2016]. For practical considerations concerning probe set design, we refer the reader to work by Cohen et al. [2018].

Test samples consisted of biofilms that were collected from the dorsum of the tongue. Samples were collected by gently scraping with a ridged plastic strip along the surface of the tongue of healthy human volunteers recruited and sampled according to a study protocol and informed consent document approved by the Institutional Review Board of the Forsyth Institute, Cambridge, MA. Samples were fixed in 50% ethanol and hybridized using a set of probes specific to different taxa of bacteria (see Table I). Two samples were taken from each of five human subjects (A-E). An additional biofilm sample was collected from subject D, chemically fixed, and hybridized without a fluorophore to generate a no-probe control sample (used to measure autofluorescence). After hybridization, the reference samples, test samples, and no-probe control were mounted on slides as described by Mark Welch et al. [2016].

**TABLE I.** OLIGONUCLEOTIDE PROBES AND THEIR TAXONOMIC TARGETS

| Probe ID | Taxon | Target | Fluorophore |
| --- | --- | --- | --- |
| Smit651-DY415-2 | Species | *Streptococcus mitis* | DY-415 |
| Ssal372-DY490-2 | Species | *Streptococcus salivarius* | DY-490 |
| Prv392-AT520-2 | Genus | *Prevotella* | ATTO 520 |
| Vei488-AT550-1 | Genus | *Veillonella* | ATTO 550 |
| Act118-TRX-1 | Genus | *Actinomyces* | Texas Red-X |
| Nei1030-AT620-2 | Family | *Neisseriaceae* | ATTO 620 |
| Rot491-AT655-2 | Genus | *Rothia* | ATTO 655 |

### B. Imaging and preprocessing

All spectral micrographs were acquired on a Zeiss LSM 880 using a 63×/1.4NA Plan-Apochromat objective. Point scanning was performed simultaneously with the 405 nm, 488 nm, 561 nm, and 633 nm lasers using the 405 and 488/561/633 dichroic mirrors. Spectral data was collected from a range of 410 nm to 696 nm using 8.6 nm steps. Each reference image was acquired using dimensions of 512 × 512 pixels at 0.263 *μ*m/pixel resolution. Test images were were acquired at 2048 × 2048 pixels and down-sampled to 1024 × 1024 pixels at a resolution of 0.220 *μ*m/pixel.

The use of dichroic mirrors blocked the detection of emitted light near the excitation wavelengths. Since these dark bands contained little information, they were removed from the reference and test images prior to any analysis. In total, 6 of the 32 spectral bands were removed.

### C. Affine mixing model

The vast majority of spectral unmixing methods assume that the emitted light from different fluorophores combines according to the linear mixing model (LMM) described in Eq. 2. The LMM is so widely used that the phrase “linear unmixing” is regularly used interchangeable with “spectral unmixing.” As point scanning confocal microscopy data is predominately read-noise-limited [Lambert and Waters, 2014], the noise component is expected to follow a Gaussian distribution with zero mean. In reality, nearly all microscopy images are contaminated by background fluorescence that offsets the noise profile [Waters, 2009, Waters and Wittmann, 2014] and breaks the assumption of zero-centered noise. To explicitly accommodate for the presence of background fluorescence, we define an affine mixing model (AMM) as

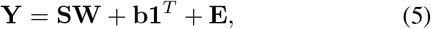

where 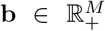 is the nonnegative background spectrum and **1** is a vector of ones. Conveniently, the AMM becomes equivalent to the LMM when no background fluorescence exists (i.e., **b** = **0**).

### D. Estimating endmembers

Spectral unmixing by NLS carries the assumption that the endmember spectra of both fluorophores and autofluorescence are known in advance. Our proposed unmixing method relaxes this assumption by only requiring that the endmember spectra of the *K* fluorophores be known. This claim is valid because one can control which fluorophores are used to label the sample, and endmember spectra can be estimated from reference images. But how to estimate endmembers from their corresponding reference images is not widely discussed in the literature. This open problem is important to emphasize because even slight inaccuracies in endmember estimates are known to dramatically effect the determination of the endmember weights—a problem called proportion indeterminacy [Zare and Ho, 2014]. We address this open question by proposing an affine nonnegative matrix factorization (ANMF) method for estimating endmembers.

For the following discussion of endmember estimation, we will denote the set of reference images as {**R**_1_, …, **R**_*K*_} with 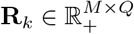, where *Q* is the number of pixels in the reference image. The corresponding endmembers, denoted **s**_*k*_ for *k* = 1, …, *K*, make up the columns of the fluorophore endmember matrix.

#### 1) Mean estimated endmember

Spectral microscopy practitioners estimate endmembers from reference images using a variety of methods based on the arithmetic mean. In general, endmembers are determined by the average spectral signature over all foreground pixels or some user-defined region of interest within the reference image. Since the illumination conditions can vary between reference and test images, it is common to normalize the mean endmembers by their *ℓ*^∞^- or *ℓ*^1^-norm. We can write this estimation procedure in matrix form as

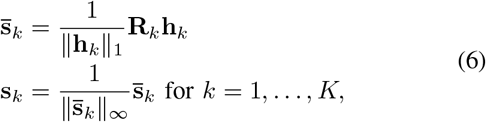

where **h**_*k*_ ∈ ℝ^*Q*^ is a binary vector with 1 indicating the foreground and 0 indicating the background of the *k*^*th*^ reference image. Foreground/background thresholding can be performed using any number of different thresholding algorithms. The Triangle algorithm is used in this work for its robustness to different illumination conditions [Zack et al., 1977].

#### 2) ANMF estimated endmember

Using the arithmetic mean of foreground pixels assumes that the noise is Gaussian distributed with zero mean. Yet, even reference images contain some level of background fluorescence. We, therefore, define an affine model similar to Eq. 5 for reference images as

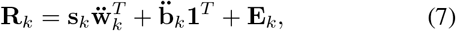

where 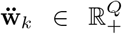 and 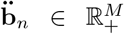 are the weights and background spectrum for the *k*^*th*^ reference image, respectively. From Eq. 7, we can formulate an endmember estimation method based on ANMF. For each fluorophore endmember spectrum 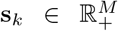 from *k* = 1, …, *K* we solve the constrained optimization problem

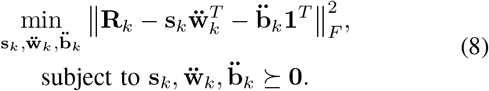

Since **s**_*k*_, 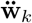, and 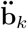 are all unknown, Eq. 8 is a nonconvex optimization problem. We also note that the solutions for **s**_*k*_ and 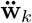 are non-unique as they may be scaled by a nonzero constant (i.e., 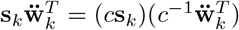). To control the uniqueness of the solution, we normalize **s**_*k*_ to the *ℓ*^∞^-norm at each iteration of the optimization scheme. This normalization is also required later to properly scale the unmixed image to the bit-depth of the observed image.

Since ANMF operates on the entire image, there is no need to threshold foreground from background or define a region of interest as with the Mean method. This is important to emphasize because a Mean estimated spectrum can change dramatically from one thresholding method to another.

### E. Semi-blind sparse affine spectral unmixing

Although it is possible to accurately characterize endmembers for each fluorophore, estimating autofluorescence endmembers is difficult for several reasons. First, it may not always be possible to create a reference sample for each type of autofluorescence that occurs in the test samples. Second, the structure of the autofluorescent tissue may influence its spectrum (i.e., thicker samples may induce more light scattering). Therefore, we believe it is better to learn the *L* distinct autofluorescence endmembers directly from the test images. This approach provides the flexibility necessary to adjust to varying numbers and types of autofluorescence spectra on an image-by-image basis.

Since we are confident in the estimation of our *K* fluorophore endmembers, it would not make sense to impose the same flexibility on them. In fact, this flexibility can cause problems during unmixing. Although a sample can be labeled with a set of fluorophores, there is no guarantee that all *K* fluorophores will exist in every field-of-view. In this rank-deficient case (i.e., *rank*(**Y**) < *rank*(**S**)), the columns of **S** that are associated with the missing fluorophores will learn some other spectrum from the image that may not be physically meaningful. Therefore, we propose a semi-blind unmixing that separates the endmember matrix **S** into two pieces: one for the *K* fluorophore endmembers, denoted 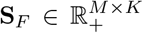 and the other for the *L* autofluorescence endmembers, denoted 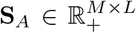. Note that the total number of endmembers is equal to the number of fluorophore and autofluorescent endmembers (i.e., *N* = *K* + *L*). To further reduce overfitting and protect against linearly dependent endmembers, we include a term to enforce sparsity of the weight matrix. We define our semi-blind sparse affine spectral unmixing (SSASU) method as

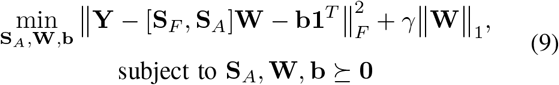

where [**S**_*F*_, **S**_*A*_] = **S** is the partitioned endmember matrix, ‖·‖_1_ is the sum over all matrix entries, and *γ* is the parameter controlling sparsity.

**Algorithm 1:**
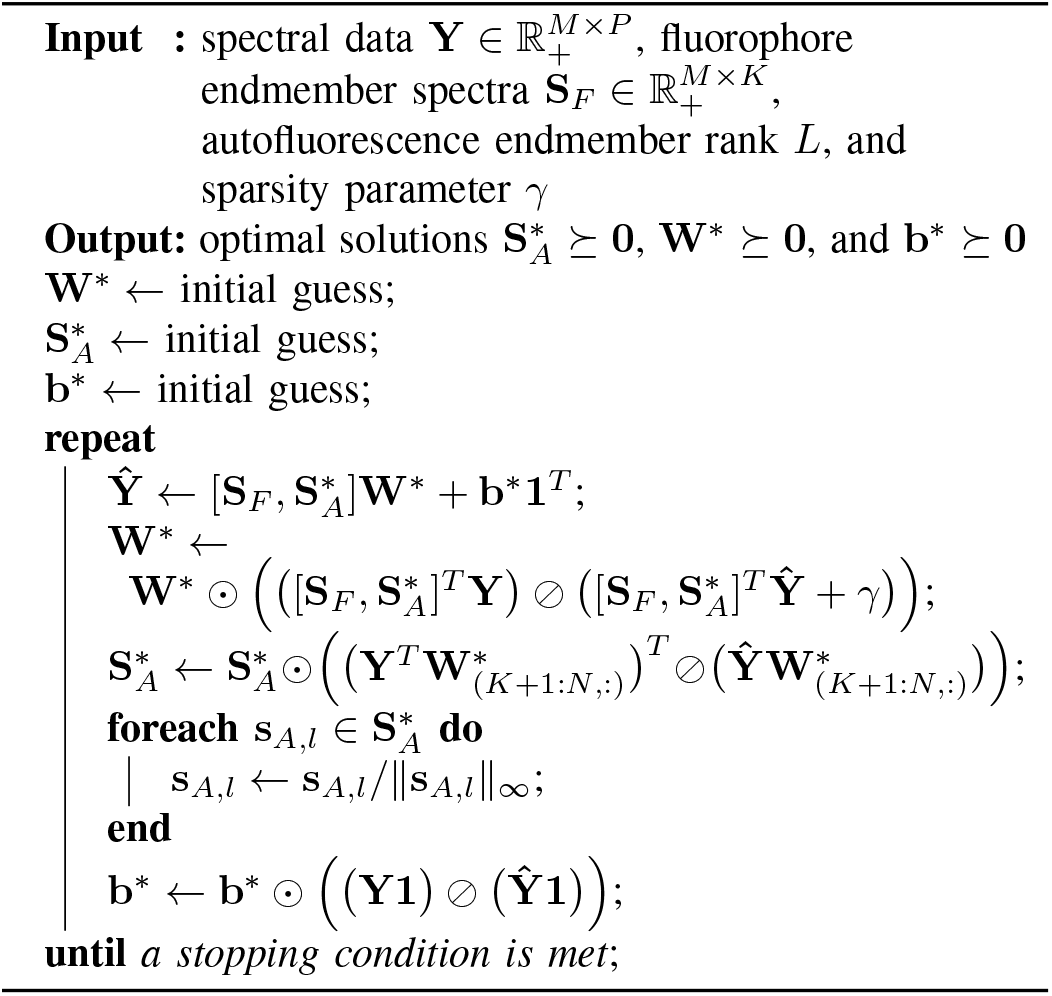
SSASU

As with our ANMF method for estimating endmembers, SSASU is a nonconvex optimization problem. However, we note that SSASU is convex in **S**_*A*_ when **W** and **b** are fixed, convex in **W** when **S**_*A*_ and **b** are fixed, and convex in **b** when **S**_*A*_ and **W** are fixed. Therefore, SSASU can be solved using a block coordinate descent algorithm similar to that suggested by Cichocki et al. [2009]. Algorithm 1 describes the multiplicative update method used to solve Eq. 9. Note that element-wise multiplication and division are denoted by ⊙ and ⊘, respectively. As with all multiplicative update algorithms, SSASU is sensitive to values becoming zero during the optimization. Therefore, we set all non-positive entries to some small value before each iteration. These small values were removed by thresholding after reaching a stopping condition.

## III. RESULTS

We assessed the robustness of endmember estimation by ANMF and arithmetic mean across seven different reference images: DY-415, DY-490, ATTO 520, ATTO 550, Texas Red-X, ATTO 620, and ATTO 655. Each endmember estimate was compared against a ground truth endmember by measuring the spectral angle.

We evaluated our SSASU method by unmixing a set of ten real-world spectral images each of which were labeled with seven fluorophores (see Table I) and contaminated by background and autofluorescence. For comparison, we performed the same evaluation with the four most commonly used unmixing methods—NLS, PoissonNMF, SUnSAL, and SUnSAL-TV. The success of each method was measured by its ability to reduce proportion indeterminacy for each test image.

### A. Endmember estimation performance

Since poorly estimated endmembers can degrade the overall performance of unmixing, it remains important to evaluate the accuracy of the estimates. Yet, defining a ground truth set of endmembers is difficult because all images contain some level of background fluorescence. Instead, we compared the endmember estimates to fluorometer data reported in the literature [McNamara et al., 2006]. Although the fluorometer data was measured under different optical and environmental conditions, these data still provided a useful baseline to check estimates made from reference images.

While neither the Mean nor the ANMF estimation method perfectly matched the fluorometer data due to differing environmental conditions, excitation wavelengths, presence of dark bands, and spectral accuracy [Cole et al., 2013], Figure 1 shows that ANMF endmembers were less contaminated by background fluorescence than Mean endmembers. We evaluated this quantitatively by calculating the spectral angle between each Mean and ANMF endmember and its corresponding fluorometer-measured spectrum. The spectral angle, which is related to cosine similarity, was calculated as 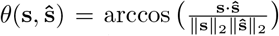, where **s** was the fluorometer spectrum and 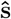 was the estimated spectrum. As reported in Table II, ANMF endmembers were as good or better at approximating the true endmember spectra than estimation by arithmetic mean.

**Fig. 1.**
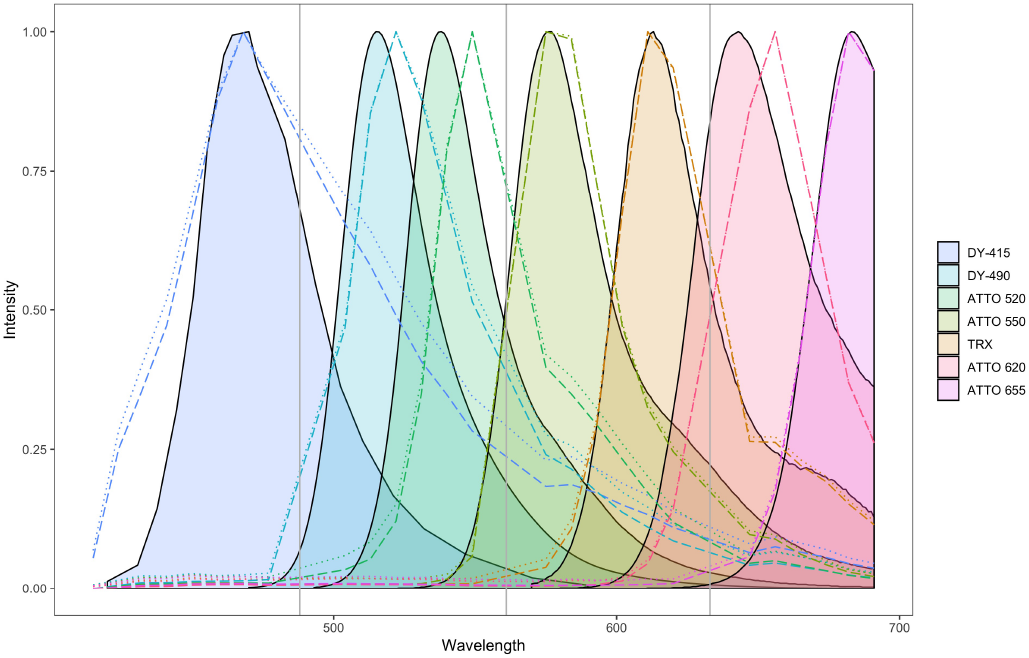
Comparison of seven Mean and ANMF estimated endmember spectra with fluorometer measurements. The shaded regions represent the fluorometer data, the dotted lines represent the Mean estimates, and the dashed lines represent the ANMF estimates. The gray vertical lines show the wavelength where dichroic mirrors blocked the measurement of emitted light (i.e., locations of missing spectral data). Note that the missing spectral data causes some endmember estimates to deviate from the fluorometer measurements (e.g., ATTO 620).

**TABLE II.** SPECTRAL ANGLE BETWEEN ESTIMATED ENDMEMBERS AND THEIR CORRESPONDING FLUOROMETER SPECTRA

| Method | DY-415 | DY-490 | ATTO 520 | ATTO 550 | TRX | ATTO 620 | ATTO 655 |
| --- | --- | --- | --- | --- | --- | --- | --- |
| Mean | 0.484 | 0.324 | 0.455 | 0.088 | 0.065 | 0.203 | 0.053 |
| ANMF | 0.421 | 0.293 | 0.455 | 0.087 | 0.052 | 0.203 | 0.024 |

### B. Parameters

SSASU, PoissonNMF, SUnSAL, and SUnSAL-TV required regularization parameters to be set prior to unmixing. Proper selection of these parameters was critical to ensuring a fair comparison of proportion indeterminacy. Therefore, parameters for SSASU, PoissonNMF, SUnSAL, and SUnSAL-TV were determined such that their unmixing solutions produced relative reconstruction errors (i.e., ±0.05) similar to NLS. Parameters were kept constant across all test images to show the flexibility of each algorithm. Both SSASU and PoissonNMF required two parameters: one controlling the sparsity of the weight matrix and the other estimating the rank of the autofluorescence endmember matrix (i.e, *L* ≈ *rank*(**S**_*A*_)). In practice, *L* is determined by manually examining the spectra of contaminated image regions. SUnSAL only needed a sparsity parameter; whereas, SUnSAL-TV required a sparsity parameter and a spatial TV parameter. The empirically determined regularization parameters used for all test images are listed in Table III. The fluorophore endmembers (i.e., **S**_*F*_) used for SSASU, NLS, PoissonNMF, SUnSAL, and SUnSAL-TV were set to the values estimated by the ANMF method. The autofluorescence endmember for NLS, SUnSAL, and SUnSAL-TV were estimated by ANMF from a no-probe control sample (i.e., an unlabeled sample taken from the dorsum of the tongue).

**TABLE III.** EMPIRICALLY DETERMINED REGULARIZATION PARAMETERS FOR EACH UNMIXING ALGORITHM

| Parameter | SSASU | PoissonNMF | SUnSAL | SUnSAL-TV |
| --- | --- | --- | --- | --- |
| No. Autofluor. Endmembers | 1 | 1 | - | - |
| Sparsity | 0.009 | 6 | 0.005 | 0.005 |
| Total Variation | - | - | - | 0.005 |

### C. Unmixing performance

The unmixing performance of SSASU, NLS, PoissonNMF, SUnSAL, and SUnSAL-TV was evaluated on the basis of two relative error metrics that ranged from zero (better) to one (worse). The ability to reconstruct the observed signal was determined by the relative reconstruction error (RRE), and was defined by

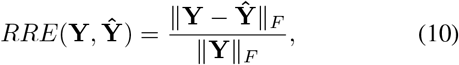

where **Y** was the observed image and 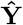 was the reconstructed image. The RRE was used to determine the parameters for each algorithm and ensure that each method properly accounted for the observed signal. It is worth noting that the RRE provides a measure of underfitting/overfitting and does not indicate the quality of a solution. For example, an algorithm can produce a physically meaningless solution to the unmixing problem with a near-zero RRE by fitting the noise in addition to the signal. Therefore, we use the RRE in conjunction with a metric that evaluates the ability of the unmixing algorithm to separate autofluorescence from the fluorophore endmembers.

Since the fluorophores used to label the test images were known to bind to distinct bacteria, each pixel in the test images contained at most one fluorophore (with the rare exception of areas where different bacteria overlap). In this case, a poor solution for an observed spectrum would produce positive weights for more than one fluorophore endmember. This is a type of overfitting known as proportion indeterminacy (PI). We measured PI by checking the non-orthogonality of the weight matrix. Since each pixel should contain only one endmember, the property **WW**^*T*^ = **D** holds, where 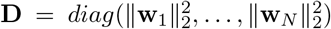 and **w**_*n*_ is the *n*^*th*^ row of **W**. From this property, we defined a measure of PI as

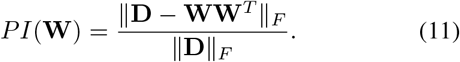

Since PI measures the intensity overlap between channels of an unmixed image, this metric would not be appropriate for ratiometric applications (e.g., FRET). For the present application, however, RRE and PI effectively indicate the fit and quality of each unmixing solution.

SSASU, NLS, PoissonNMF, SUnSAL, and SUnSAL-TV were each able to effectively reconstruct the test images with RREs below 0.15 and within 0.05 of each other (see top of Figure 2). Across all test images, SUnSAL-TV had the highest RRE. This was expected because SUnSAL-TV is the only algorithm with both a sparsity and a spatial regularizer. The consistent RREs across all images suggests that none of the methods significantly overfit or underfit the observed data.

**Fig. 2.**
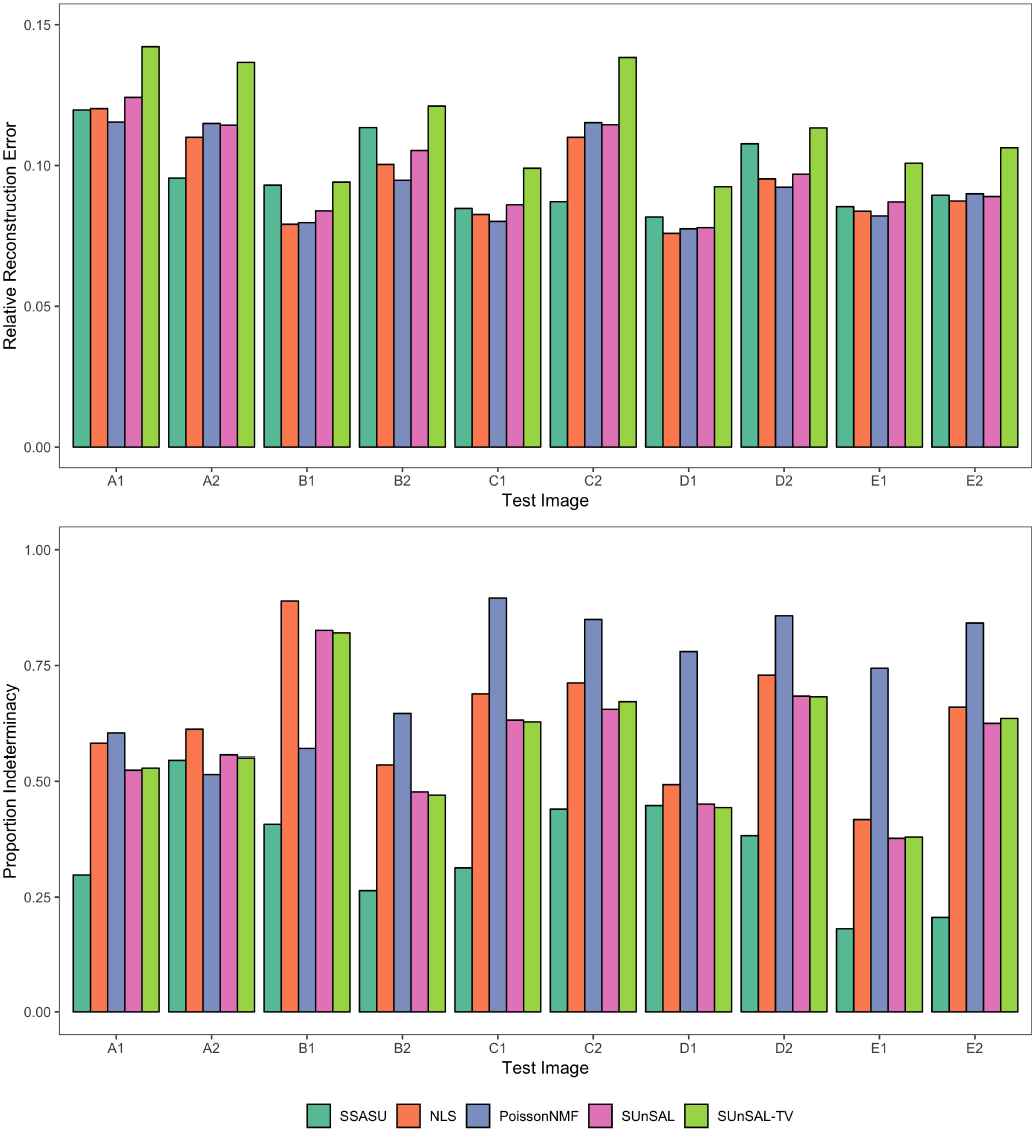
Comparison of unmixing performance for SSASU, NLS, and PoissonNMF across ten test images taken from five samples. The relative reconstruction error (top) evaluates each method’s ability to reconstruct the observed spectra image. The proportion indeterminacy (bottom) measures the non-orthogonality of the weight matrices and illustrates how well each method separates the fluorophore endmembers in the presence of autofluorescence. Both metrics range from zero (better) to one (worse).

Despite having similar RRE across all test images, the quality of the solutions varied dramatically between the different methods. SSASU outperformed NLS, PoissonNMF, SUnSAL, and SUnSAL-TV in all but two test cases at reducing PI (see bottom of Figure 2). SSASU performed second best in PI for the remaining two test images. For eight of the ten test images, PoissonNMF was least able to cleanly separate the fluorophores. Although there was no noticeable difference in PI between SUnSAL and SUnSAL-TV, they both consistently produced better quality solutions than NLS. When comparing the worst PI for each method, SSASU showed a clear improvement over NLS, PoissonNMF, SUnSAL, and SUnSAL-TV at 0.55, 0.89, 0.90, 0.83, and 0.82, respectively.

The performance illustrated by these metrics can be observed qualitatively in the unmixing results for test image E2 (see Figure 3). In the results for NLS (Figure 3 A-H), autofluorescence has contaminated nearly all of the unmixed channels. This same autofluorescence light was efficiently captured in the autofluorescence channel of the SSASU unmixed image (Figure 3 I). We also note that *Prevotella* (ATTO 520) was not present in image E2, yet the ATTO 520 channel of the NLS unmixed image contained a significant amount of light (Figure 3 D). Using the parameter settings described in Table III, we note that SSASU was unable to remove all proportion indeterminacy found in the DY-415 and DY-490 channels as indicated by the repeated structures (Figure 3 J & K). Nevertheless, the composite view of all unmixed fluorophore channels (i.e., all channels excluding autofluorescence) clearly illustrates the ability of SSASU to efficiently separate both the autofluorescence and the fluorophores (Figure 3 Q & R).

**Fig. 3.**
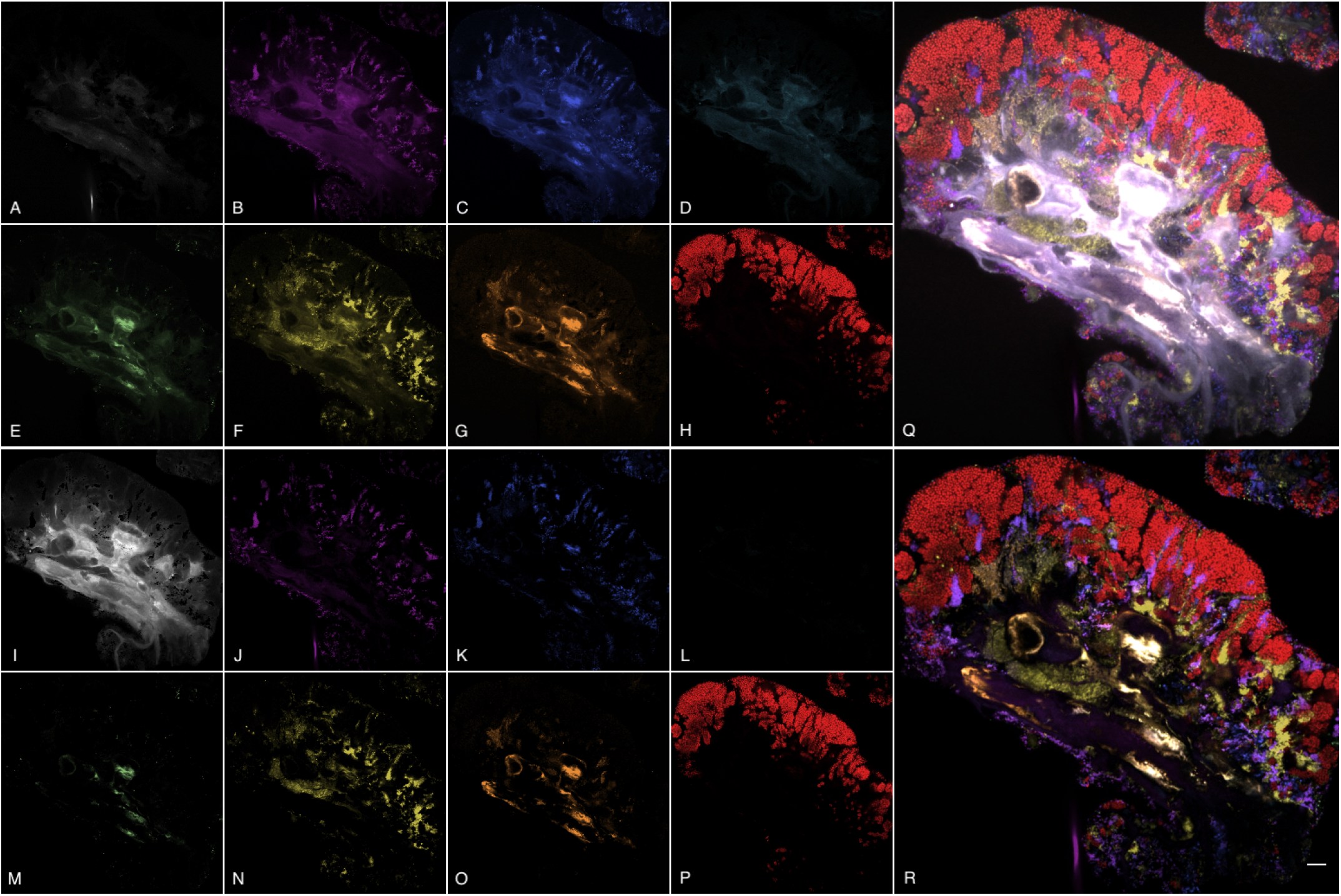
Montage of unmixed images for NLS (top) and SSASU (bottom). Panels A-P show the unmixed channels for autofluorescence (A, I); *S. mitis*/DY-415 (B, J); *S. salivarius*/DY-490 (C, K); *Prevotella*/ATTO 520 (D, L); *Veillonella*/ATTO 550 (E, M); *Actinomyces*/Texas Red-X (F, N); *Neisseriaceae*/ATTO 620 (G, O); and *Rothia*/ATTO 655 (H, P). A larger composite view of the non-autofluorescence unmixed channels is shown for NLS in panel Q and for SSASU in panel R. The scale bar in panel R indicates 10 *μ*m.

### D. Characterizing autofluorescence

Samples can contain many different types of autofluorescent molecules, and exactly which autofluorescence endmembers exist in an image is difficult to ascertain in advance. For applications using NLS, SUnSAL, or SUnSAL-TV, it is common to estimate the autofluorescence endmember from a no-probe control. Figure 4 shows how the autofluorescence endmember estimated from the no-probe control image compared to those learned by SSASU. It is clear that the no-probe control endmember poorly characterized the autofluorescence encountered in the test images. The learned autofluorescence signatures are consistent with the spectra of lipids, collagen, and other common fibrous proteins [Croce and Bottiroli, 2014]. The ability to learn the autofluorescence spectrum directly from an image allows SSASU to adapt to highly varied samples. This flexibility lets SSASU fit the observed data with a sparse set of endmember weight, thereby reducing proportion indeterminacy as compared to NLS, SUnSAL, and SUnSAL-TV.

**Fig. 4.**
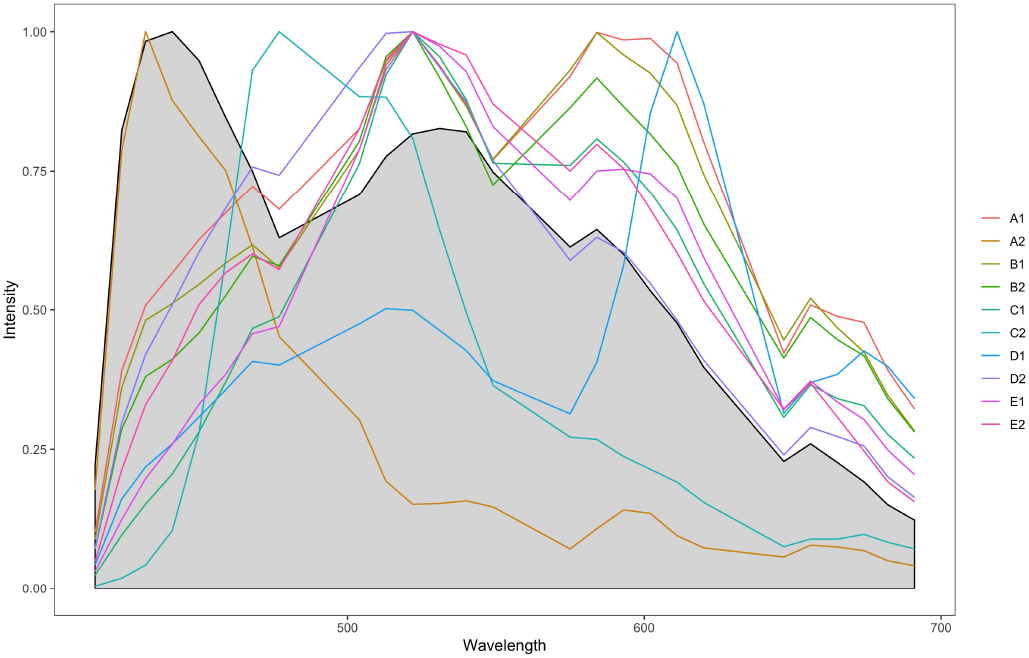
Comparison of the autofluorescence endmember estimated from the no-probe control reference image (gray region) to the autofluorescence endmembers learned by SSASU.

## IV. CONCLUSION

Spectral microscopy and unmixing make it possible to visualize biological samples labeled with a large set of fluorophores. However, the choice of unmixing algorithm is important for achieving the desired results. In this paper, we proposed and evaluated a semi-blind sparse affine spectral unmixing (SSASU) algorithm aimed at separating fluorescence endmembers in the presence of autofluorescence and background fluorescence. In all but two test cases, SSASU was able to outperform NLS, PoissonNMF, SUnSAL, and SUnSAL-TV in mitigating autofluorescence. While our method is more flexible than NLS, we note that, like other NMF methods, SSASU is more computationally expensive and does not guarantee a unique solution. Therefore, we recommend SSASU for situations where spectral micrographs are contaminated with one or many sources of autofluorescence.

In addition, we described an affine nonnegative matrix factorization (ANMF) method for estimating endmembers from reference images. We showed that ANMF estimation was as good or better than the Mean method across all test cases. In addition, ANMF does not depend on a thresholding of foreground and background. This makes ANMF more robust to images with uneven illumination profiles.

There are several obvious extensions of this work. First, formulating a version of SSASU for tensors would allow for the unmixing of spectral images that use sequential excitation. Second, allowing minor adjustments to fluorophore endmember spectra on an image-by-image basis would allow the algorithm to accommodate endmember variability (i.e., changes in the endmember spectra as a result of the microenvironment).

## ACKNOWLEDGEMENTS

The authors would like to thank Andrew Kempchinsky for his help with sample preparation; Dr. Lars Ruthotto, Dr. Yuanzhe Xi, Yunyi Hu, and Kelvin Kan for their thoughtful advice and guidance; and Louis Kerr for his microscopy support.

